# IgMAT: immunoglobulin sequence multi-species annotation tool for any species including those with incomplete antibody annotation or unusual characteristics

**DOI:** 10.1101/2021.09.22.461368

**Authors:** Daniel Dorey-Robinson, Giuseppe Maccari, Richard Borne, John A. Hammond

**Affiliations:** The Pirbright Institute, Pirbright, United Kingdom; Anthony Nolan Research Institute, London, United Kingdom

## Abstract

The advent and continual improvement of high-throughput sequencing technologies has made immunoglobulin repertoire sequencing accessible and informative regardless of study species. However, to fully map changes in polyclonal dynamics, precise annotation of these constantly rearranging genes is pivotal. For this reason, data agnostic tools able to learn from presented data are required. Most sequence annotation tools are designed primarily for use with human and mouse antibody sequences which use databases with fixed species lists, applying very specific assumptions which select against unique structural characteristics. We present IgMAT, which utilises a reduced amino acid alphabet, incorporates multiple HMM alignments into a single consensus and enables the incorporation of user defined databases to better represent their species of interest.

**Availability and implementation:** IgMAT has been developed as a python module, and is available on GitHub (https://github.com/TPI-Immunogenetics/igmat) for download under GPLv3 license.

**Supplementary information:** Model Breakdowns

## INTRODUCTION

Whole antibody repertoire sequencing can result in millions of sequences which can require various layer of filtering for specificity and quality. Perhaps the most important step is accurate annotation of the framework (FR1-4) and complementary determining (CDR1-3) regions that underpins the accuracy of most downstream analyses. Whilst numerous web servers exist with such functionality, the ability to run a tool locally as part of in-house workflows is required for many projects. One such tool is ANARCI (Dunbar and Deane, 2015), which applies a range of numbering schemes to annotate input sequences by applying Hidden Markov Models (HMMs) trained with curated data from the IMGT database (Giudicelli et al., 2005). However, this approach does not provide adequate flexibility to annotate antibody sequences from species with unusual structural properties. This is problematic where species have incomplete IMGT records or use genetic mechanisms that may inhibit alignments to standard gene sequences such as gene conversion in chickens (Arakawa et al., 2002). Further, tools designed using assumptions based on model species such as human and mouse inefficiently capture or exclude unusual antibodies, for example imposing CDRH3 maximum length fails to identify ultralong antibodies in cattle [Deiss et al., 2019].

Here we present IgMAT (Immunoglobulin Multispecies Annotation Tool), a tool for the automatic discrimination and annotation of antibody sequences, specifically designed to be integrated into custom analysis pipelines. IgMAT is based on the ANARCI tool, with extended capability to annotate antibody sequences from multiple species. The tool is highly customizable, allowing the addition of custom antibody sequence datasets and generating a range of output formats including a bed file of FR and CDR coordinates, enabling downstream analyses as required.

## THE IgMAT PACKAGE

IgMAT provides convenient tools for the analysis and annotation of antibody sequences, allowing the analysis of millions of sequences at the same time. Like many other antibody numbering tools (ANARCI, PyIgClassify, ProABC (Adolf-Bryfogle et al., 2015; Dunbar and Deane, 2015; Olimpieri et al., 2013)), the algorithm applies a set of precomputed HMMs to align the input sequences according to the IMGT numbering schemes (Lefranc et al. 2003) and successively perform annotation. IgMAT uses a dataset of curated germline antibody sequences of different domains for a set of organisms from the IMGT/Gene Database (Giudicelli et al., 2005). Additionally, the use of custom datasets allows IgMAT to include unusual antibody sequences. This can be extremely useful for annotating sequences with unusual length or recombination patterns.

## ALGORITHM

IgMAT can annotate single sequences or batches efficiently by distributing jobs among multiple processes. Each sequence is aligned to the HMMs to find the best matching domain. For the most common antibody sequences, one single match is sufficient to identify all the regions composing the antibody sequence (FR1-4, CDR1-3). However, some antibodies can display ultralong CDR3 sequences or unusual patterns that are not identified by one single HMM match. For this reason, IgMAT considers multiple HMM alignments from the same domain and extracts a consensus sequence that is then validated by applying heuristic knowledge of FR and CDR regions derived from the input model. This approach allows annotation of most known antibody sequences. However, it is limited by the number and variability of the sequences composing the input dataset, and for some extreme cases it cannot guarantee a proper annotation. To overcome this limitation, IgMAT implements two additional features: a tool for generating custom HMM models and the ability to use a reduced amino acid alphabet.

IgMAT employs a data extraction script based on the code from ANARCI to generate the default HMM model with the high-quality germline sequences from the IMGT. In addition, IgMAT provides the ability to build custom HMM datasets that can be used to annotate sequences. The build tool requires an initial set of FASTA files containing J region and V region sequences separately. An alignment is then automatically generated from all permutations of the VJ sequences; after validation of the alignment file, the HMM is created and a name is assigned to the model. Optionally, the script can directly accept an alignment file to allow the user to fix any possible annotation errors before the HMM model is created. The ability of IgMAT in recognising and annotating antibody sequences is directly correlated to the coverage provided by the input dataset. If an input sequence is not represented by the dataset, it won’t be recognised by the HMM. Reducing the number of amino acids composing the sequence helps simplify the input data, highlighting chemo-physical patterns that are not properly represented by the input dataset. IgMAT implements the reduced alphabets by Li et al (Li et al., 2003), providing the ability to apply a reduction from twenty amino acids down to three.

## MULTISPECIES TESTING

IgMAT functionality and versatility was tested by analysing a panel of high-throughput sequencing data obtained from a variety of technologies and species. Whole repertoire data from horse (Equus caballus) (Manso et al., 2019), mouse (Mus musculus) (Rettig et al., 2018), camel (Camelus bactrianus) (Li et al., 2016), human (Homo sapiens) (Galson et al., 2020), pig (Sus scrofa) (Schwartz. 2013) and chicken (Gallus gallus), as well as multiple combined single cell datasets from cattle (Bos taurus) (Li et al., 2019) were annotated using IgMAT (Figure 1A, Table 1). When the species was included in the default IgMAT reference model (Table S1), or not available from IMGT the default HMM model was used, otherwise a species-specific HMM model was constructed with IMGT V and J sequences. Input sequences were translated into all six reading frames and any sequences containing a stop codon were removed. Over 80% of bovine, camelid, porcine and chicken (except chicken IgA) input sequences were successfully annotated. Horse, mouse, and human annotation rates were similar to those previously found with differences likely due to species-specific fine-tuned configuration in the analysis pipeline (Figure 1B). Where annotation failed, a large proportion of sequences were too short (15-20 amino acids) and thus failed alignment. Overall, IgMAT was able to annotate the overwhelming majority of correct antibody sequences from high-throughput sequencing data of a range of vertebrate species without having to apply tailor-made datasets.

**Table 1.**
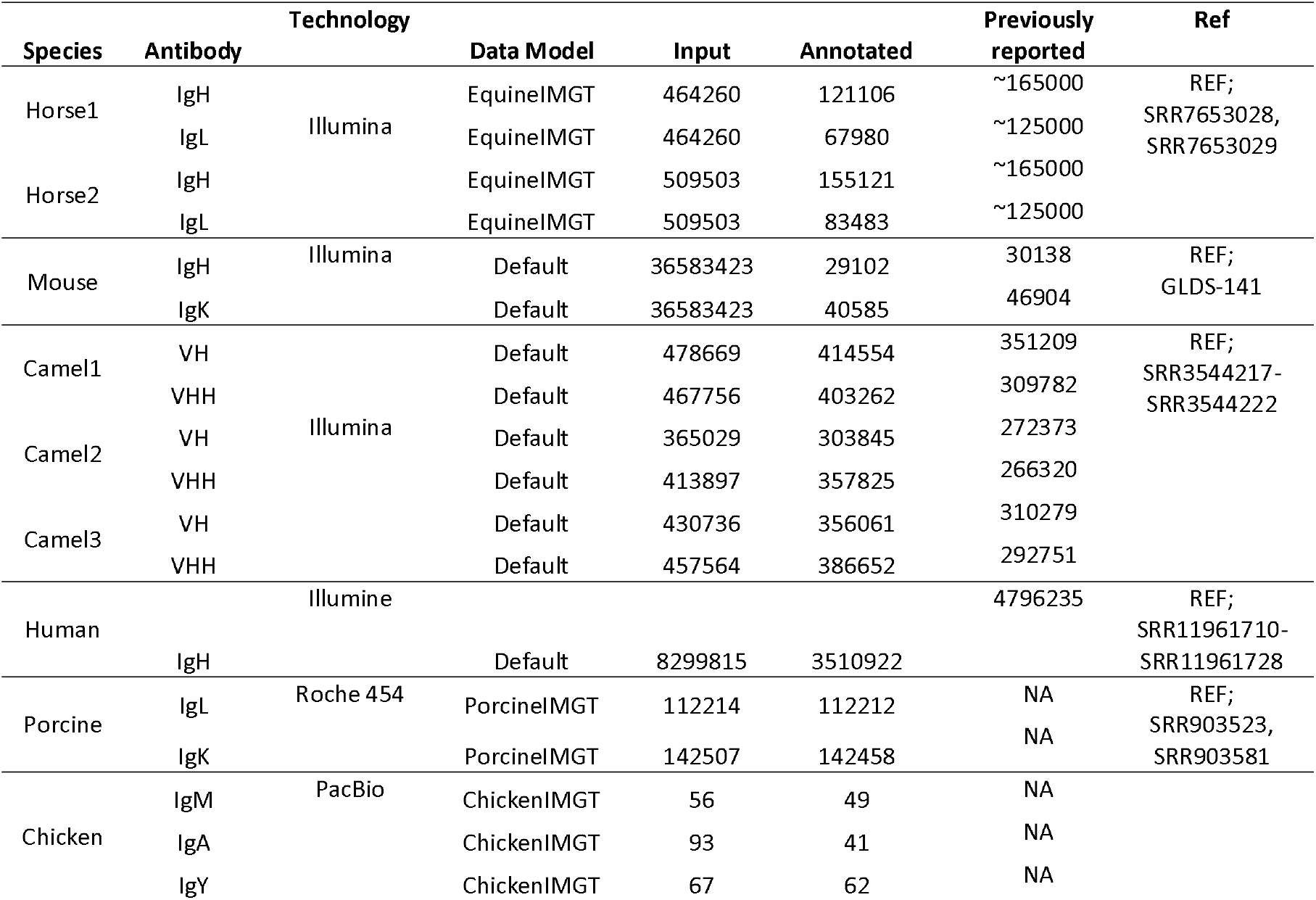
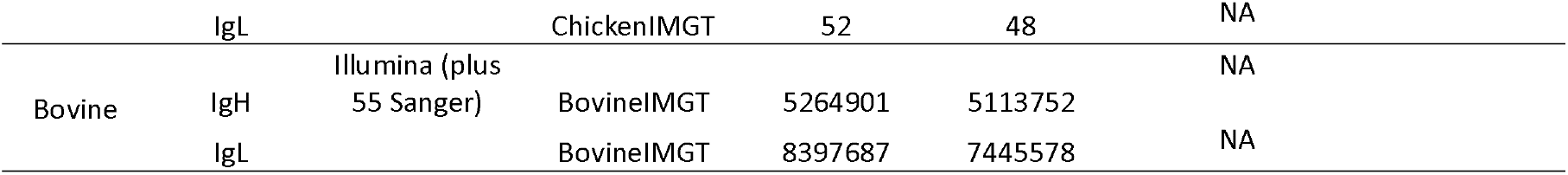
Benchmark of IgMAT annotation. Whole repertoire data from different species were analysed and annotated with IgMAT. Depending on the species, different data models were used. Th default dataset includes human, mouse, rhesus monkey, rabbit, sheep, alpaca, rat and pig IMGT sequences.

**Figure 1.**
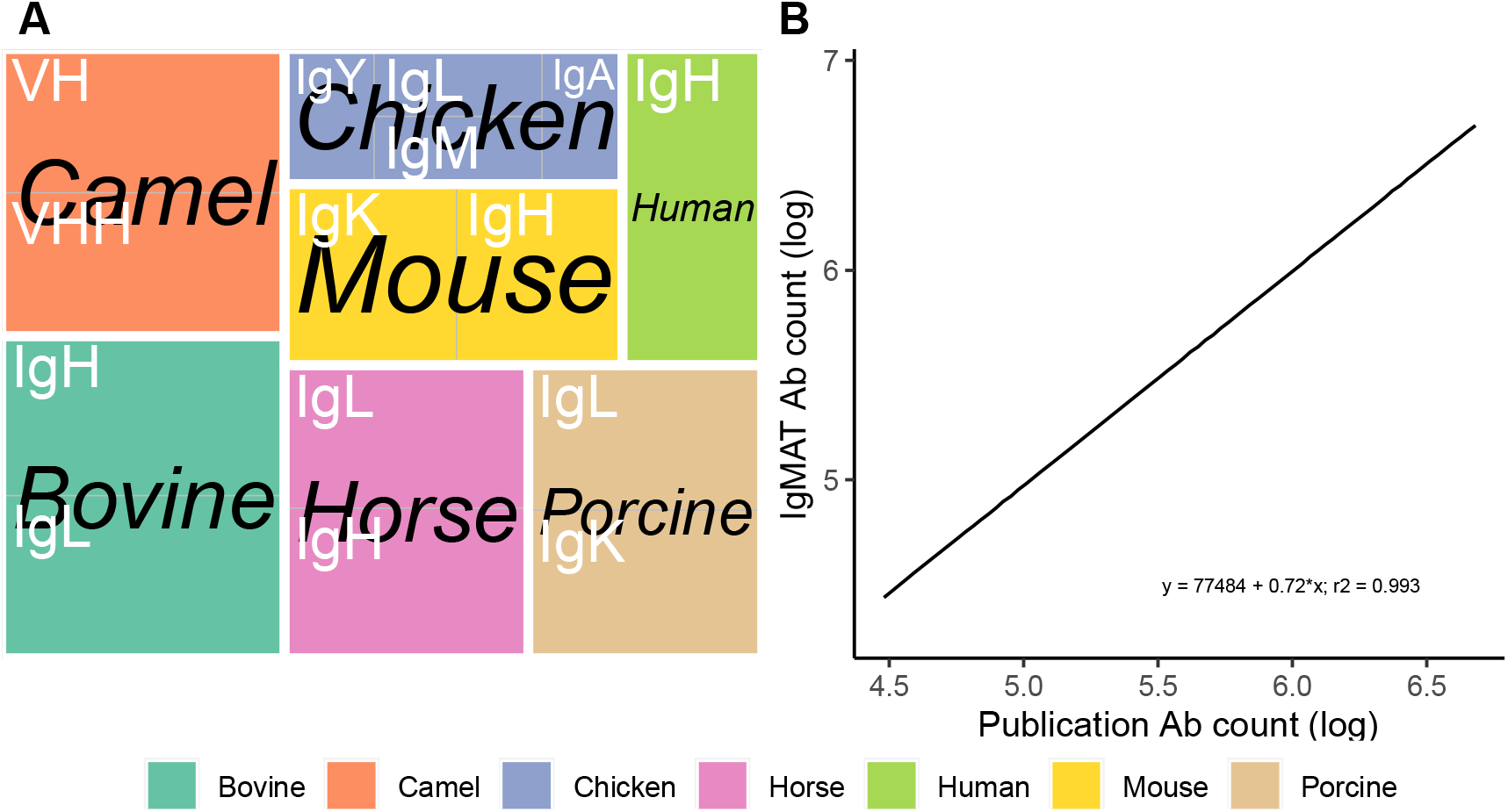
Dataset composition and evaluation results. A) The dataset used for testing comprised sequences from different species and isotypes; B) Comparison of the number of IgMAT annotated antibodies with the antibodies identified in the test dataset.

## CONCLUSION

IgMAT provides enormous flexibility to define custom models incorporating user defined data to better explore antibody repertoires from any vertebrate species, using species specific or multi species databases. Here we have demonstrated an ability to identify and annotate antibody sequences from seven species using only IMGT sequences to build the HMM data set. The addition of sequences from specialist or in house data increases the power to detect the antibody sequences of any species of interest. The underlying principle of IgMAT allows it to be readily applied to T cell receptor datasets.

## Acknowledgements

This research was supported by Bill & Melinda Gates Foundation awards OPP1192002 and OPP1215550. JAH was also supported by the United Kingdom Research and Innovation Biotechnology and Biological Sciences Research Council (UKRI-BBSRC) awards, BBS/E/I/00007030, BBS/E/I/00007031, BBS/E/I/00007038 and BBS/E/I/00007039.

